# Are methylation beta-values simplex distributed?

**DOI:** 10.1101/753459

**Authors:** Lara Nonell, Juan R González

## Abstract

DNA methylation plays an important role in the development and progression of disease. Beta-values are the standard methylation measures. Different statistical methods have been proposed to assess differences in methylation between conditions. However, most of them do not completely account for the distribution of beta-values. The simplex distribution can accommodate beta-values data. We hypothesize that simplex is a quite flexible distribution which is able to model methylation data.

To test our hypothesis, we conducted several analyses using four real data sets obtained from microarrays and sequencing technologies. Standard data distributions were studied and modelled in comparison to the simplex. Besides, some simulations were conducted in different scenarios encompassing several distribution assumptions, regression models and sample sizes. Finally, we compared DNA methylation between females and males in order to benchmark the assessed methodologies under different scenarios.

According to the results obtained by the simulations and real data analyses, DNA methylation data are concordant with the simplex distribution in many situations. Simplex regression models work well in small sample size data sets. However, when sample size increases, other models such as the beta regression or even the linear regression can be employed to assess group comparisons and obtain unbiased results. Based on these results, we can provide some practical recommendations when analyzing methylation data: 1) use data sets of at least 10 samples per studied condition for microarray data sets or 30 in NGS data sets, 2) apply a simplex or beta regression model for microarray data, 3) apply a linear model in any other case.

## Background

Epigenetic events are those that, without changing the original DNA structure, alter gene expression levels. DNA methylation, the most studied epigenetic modification, involves the addition of a methyl group to the fifth carbon of cytosine [1]. DNA methylation predominantly occurs in CpG sites, which are DNA dinucleotides composed of a cytosine nucleotide followed by a guanine nucleotide. CpG islands are regions highly enriched in CpG sites. Changes in methylation patterns and levels are diverse across tissues and have been associated with various diseases or traits such cancer, genetic disorders [2], smoking [3] aging [4] and sex [5].

There are at present two main technologies to assess methylation levels, microarrays and next generation sequencing (NGS) [1]. In the field of microarrays, Illumina Infinium is the market leader with the EPIC array, the popular 450K and the previous 27K arrays. Methylation levels obtained from microarrays are represented in terms of beta-values, which measure the proportion of methylated probes (i.e. values between 0 and 1). To assess methylation levels in NGS, methods that apply the bisulfite modification, such as whole-genome bisulfite sequencing (WGBS) and reduced representation bisulfite sequencing (RRBS), are the most demanded. Methylated and unmethylated read counts are obtained from NGS and their ratios are equivalent to the beta-values derived from microarray. Therefore the same statistical methods can be used, although coverage depth should be taken into account [6].

Methylation differences between conditions are typically reported at differentially methylated sites (DMSs), although differentially methylated regions (DMRs), which include multiple adjacent CpG sites, can also be provided. Several methods have been described to assess DMSs. In small sample size experiments, where distribution assumptions may be inaccurate, a Fisher’s exact test is often applied. Other classical hypothesis testing methods, such as the chi-square test, regression approaches, t-test and analysis of variance; are used to identify DMSs [7, 8]. Limma [9] is also an extended method to assess DMSs using standard linear regression models. Some methods assume beta-values to follow a beta distribution [10, 11].

In the specific context of NGS, coverage depth differences in samples can generate overdispersion. This has been addressed mainly by fitting a logistic regression that accounts for coverage depth (Methylkit [12]) or, more commonly, through the beta-binomial distribution. Beta-binomial was proposed by Molaro et al. [13] as the natural count-based statistical distribution [14, 7]. Posteriorly, several methods such as DSS [15], DSS-general [16], MOABS [17], and RADmeth [14] that assume a beta-binomial were designed.

Beta-values represent proportions restricted to the [0,1] interval. They can be skewed or even bimodal, with peaks close to 0 or 1. Distributions for proportional data include the beta distribution, part of the exponential family distribution; and the simplex, which belongs to the family of dispersion models [18]. Both beta and simplex are defined in the open interval (0,1) but real data contain sometimes bounds 0 and/or 1, so the corresponding 0/1 inflated distributions might be required. Since there is currently no agreement on which distribution to apply for proportional data, although recommendations to use the beta have been described in other fields [19], we want to assess the convenience of the simplex distribution to accommodate beta-values.

Association of beta-values with phenotype can be performed through regression models. The generalized linear model (GLM), which was developed for exponential families of distributions but extended to the dispersion models [20], is the standard approach to compare the means of different groups. However, as beta-values distribution may present asymmetries, quantile regression models could be an alternative in these situations.

In this paper, we aimed to study whether simplex could be a good representation for methylation beta-values, which could be in turn fitted by a simplex regression model with the use of GLMs. In addition, we wanted to assess whether quantile regression could also be a good model to analyse beta-values by capturing differences in the extreme values of the data. For that, we conducted three different analysis strategies on four real data sets, obtained from microarrays and NGS. First, the data sets were analyzed and modelled. Second, some simulations were performed to test different scenarios encompassing several distribution assumptions, regression models and sample sizes. Finally, we performed DNA methylation comparisons between females and males in order to benchmark the assessed methodologies.

## Methods

### Data sets

Data analyses were performed using four different public data sets. These include both microarrays and NGS data. Details on selected data sets are specified below and summarized in Table 1.

**Table 1:** Data sets used in the analyses.

| Data set | Origin | Project | Platform | CpGs | Filtered | Samples |
| --- | --- | --- | --- | --- | --- | --- |
| a450k-smoking | GSE50660 | Smoking effect | Illumina Infinium HumanMethylation450 BeadChip | 485,577 probes | 125,950 | 464 |
| aEPIC-PBB | GSE116339 | Polybrominated biphenyl (PBB) exposure | Illumina Infinium MethylationEPIC BeadChip | 868,564 probes | 198,711 | 679 |
| RRBS-ES | Rnbeads.org | Ewing sarcoma | RRBS | 5,658,698 | 270,569 | 188 |
| WGBS-BLUE | Rnbeads.org | BLUEPRINT | WGBS | 28,571,135 | 204,073 | 81 |

### Cigarette Smoking effects on methylation from 464 samples by Illumina 450k array

Epigenome-wide microarray association study in peripheral-blood DNA of 464 individuals who were current (n = 22), former (n = 263) and never smokers (n = 179). This research was performed on the Infinium HumanMethylation450 BeadChip array (Illumina Inc, USA). Data were downloaded from GEO (GSE50660) [21] and preprocessed using the *minfi* R package [22] by quantile normalizing and SNP purging. This led to 381,306 CpGs distributed in the open (0,1) interval. Then, data was filtered selecting those CpGs with standard deviation (sd) > 0.03 obtaining 125,950 CpGs for further analyses. This data set will be hereafter referred to as the array 450k smoking data set (*a450k-smoking*).

### Exposure to polybrominated biphenyl of 679 samples by Illumina EPIC array

In this study, the blood DNA of 679 individuals who were exposed to polybrominated biphenyl (PBB) in the 1970’s in Michigan (USA), was interrogated with the Infinium MethylationEPIC BeadChip array (Illumina Inc, USA). The processed matrix was downloaded from GEO (GSE116339), SNPs removed (using *minfi* package) and filtered by selecting those CpGs with sd > 0.03 obtaining 198,711 CpGs for further analyses. In order to determine the effect of data normalization on the results, we performed the six available normalization methods in *minfi* package and found that shape of global distribution did not change much (Supplementary figures 1 and 2) data set as beta-values were highly correlated (range of significant correlations 0.95-0.99). This data set will be hereafter referred to as the array EPIC PBB data set (*aEPIC-PBB*).

**Figure 1:**
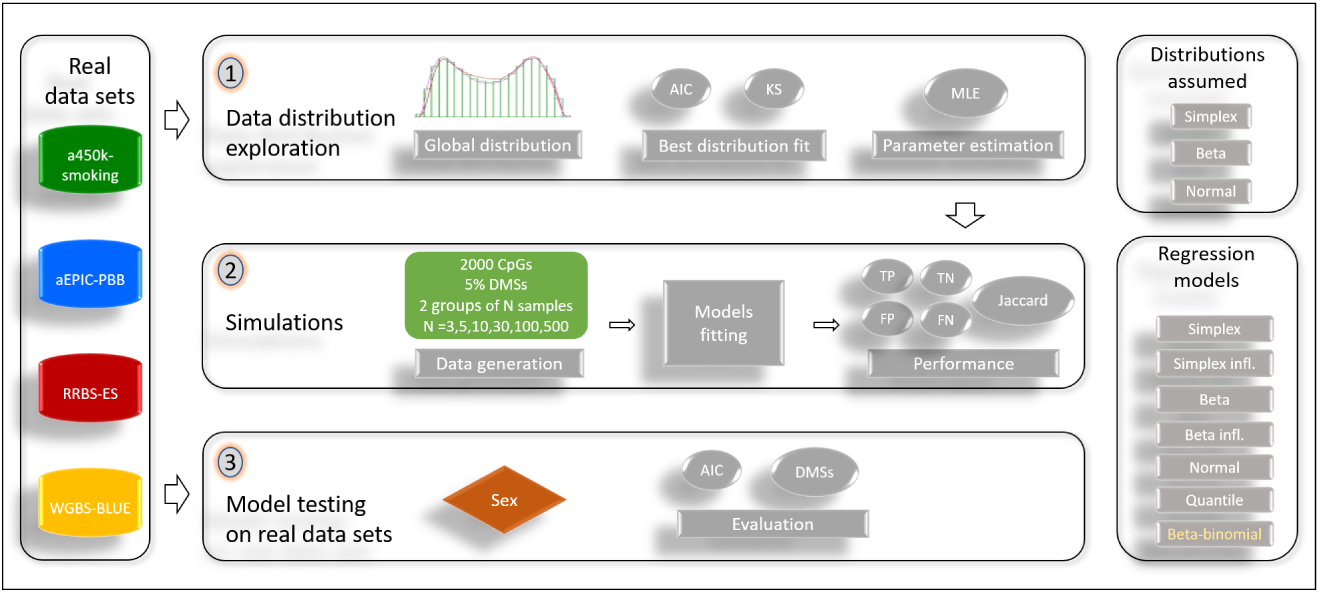
Scheme of analyses performed. AIC: Akaike’s information criterion, DMS: differentially methylated site, KS: Kolmogorov-Smirnov test, MLE: maximum likelihood estimation, a450k-smoking: array 450k smoking data set, aEPIC-PBB: array EPIC PBB data set, RRBS-ES: reduced representation bisulfite sequencing Ewing sarcoma data set, WGBS-BLUE: whole genome bisulfite sequencing BLUEPRINT data set, TN: true negatives, FP: true positives, FN: false negatives, FP: false positives.

**Figure 2:**
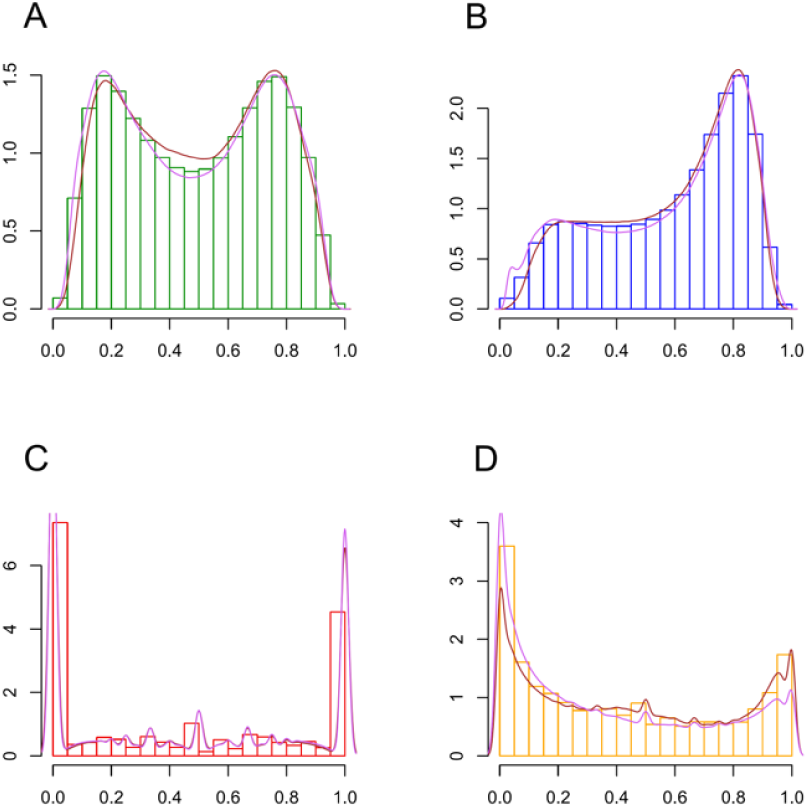
Beta-values histogram of all CpGs in the assessed data sets. Assessed conditions are added as density lines in different colors, female in brown and male in violet. A. array 450k smoking data set. B. array EPIC PBB data set. C. RRBS ES data set. D. WGBS BLUEPRINT data set.

### Genome-scale RRBS data for 188 cases suffering Ewing Sarcoma

This study assesses DNA methylation associated with Ewing sarcoma, a bone cancer primarily affecting children and young adults. In addition to tissue samples, healthy mesenchymal stem cells (MSCs), MSCs affected with Ewing sarcoma and Ewing cell lines were also included in the study [23]. Data of 188 RRBS samples were downloaded from https://rnbeads.org/methylomes.html and preprocessed using the *RnBeads* R package ([24]). From an initial number of 5,658,698 CpGs; sites with sd > 0.2 annotated to be in an *island, shelf* or *shore*; with at least 30 non-missing values and a mean coverage of 3 were retained; resulting in a total of 270,569 assessed sites. This data set will be hereafter referred to as the RRBS Ewing sarcoma data set (*RRBS-ES*).

### WGBS data of 81 blood sample methylomes from the BLUEPRINT project

Whole genome bisulfite sequencing profiles were generated for 81 different cell types obtained from blood samples of healthy donors in the BLUEPRINT project framework. Raw data were downloaded from https://rnbeads.org/methylomes.html and preprocessed using the RnBeads package. From an initial number of 28,571,135 CpGs; sites with sd > 0.2 annotated to be in an *island, shelf* or *shore* with at least 30 non-missing values and a mean coverage of 3 were conserved. Finally, a total of 204,073 sites were analysed. This data set will be hereafter referred to as the WGBS BLUEPRINT data set (*WGBS-BLUE*).

### Distributions and regression models

The main objectives of this paper are to decipher whether the real distribution of beta-values is the simplex and to identify the regression model that best fits in the association with phenotype. Table 2 summarizes the assessed distributions for methylation beta-values, their oddities and the natural regression models to assess DMSs in each particular distribution. Below we describe the beta and simplex distributions, defined on the interval (0,1), and also the regression models. The functions and R packages used in the analyses are also given.

**Table 2:**
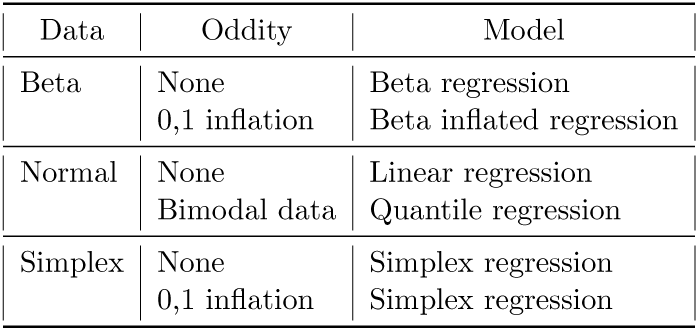
List of distributions with their oddities and their natural regression model to assess DMSs.

#### Beta distribution

The beta distribution belongs to the exponential family. The density function of the beta distribution with parameters *µ* and *ϕ* is:

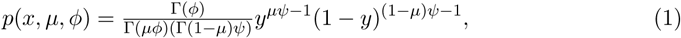

where Γ denotes the gamma function [25]. *dbeta* (package *stats*) function was used to estimate beta parameters.

#### Simplex distribution

The simplex distribution belongs to the family of dispersion models. Considering the normal distribution with mean *µ* and variance *σ*^2^ with the following density function:

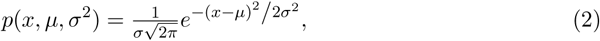

then, the simplex distribution with location parameter *µ* and dispersion parameter *σ*^2^ is defined as follows:

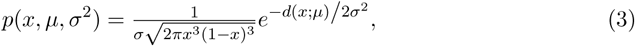

Where 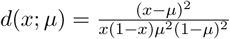 and *x* ∈ (0,1), *μ* ∈ (0,1).

Parameter estimation through the maximum likelihood method were published by Peter X.-K. Song [26]. *ZOIP* package [27] was used to estimate simplex parameters.

#### Regression model

The linear regression model has the following formulation:

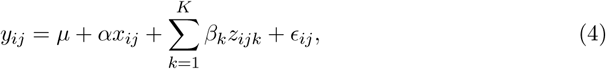

where *y* is the methylation level, *µ* is the mean of *y, x* is the phenotype or condition variable, *α* is the condition coefficient, *z* are the covariates, *β* are the estimated coefficients in the K groups and *ϵ* is the error term. *i* represents the methylation site, *j* the subject and *k* the covariates.

GLMs allow the response variable *Y* in equation (4) to adopt distributions belonging to the exponential or dispersion families distributions and can therefore be used to fit beta regression or simplex models. Beta regression was performed with *betareg* package [28] whereas 0/1 inflated beta regression was carried out with the *gamlss* package [29]. Simplex regression was fitted with *simplexreg* package [30] while inflated simplex regression was fitted with the *ZOIP* R package.

#### Quantile regression

Quantile regression was introduced by Koenker and Bassett in 1978 [31] to expand the potential of linear models. The regression coefficients are computed by minimizing the sum of weighted absolute residuals [32]. Quantile regression fits specified percentiles of the response, to accommodate the different distribution shapes and can potentially describe the entire conditional distribution of the response. 75% quantile regression was assessed for the data distribution with package *quantreg* [33].

#### Other regression models

Beta-values from microarray data were also modelled using linear models, with function *lm* from package *stats* and also *limma* package [9]. NGS data sets were analyzed using beta-binomial regression using the function *betabin* from package *aod* [34] by modelling the total methylated reads and coverage at each CpG site.

### Data analyses

We used three different approaches to compare the different data modelling strategies. Figure 1 summarizes data analyses. First, we assessed whether beta-values follow three different data distributions: simplex, beta or normal. Second, we run a comprehensive simulation study to assess and evaluate the different models under different scenarios (sample size, effect size, etc.). Finally, we tested regression models on real circumstances by comparing DNA methylation between females and males. All analyses were conducted in the R environment (v. 3.5.1). R functions used to perform all tests are available at https://github.com/lnonell/MetDist.

#### Data distribution

For each CpG, the best distribution of its beta-values across samples was assessed in terms of the Akaike’s information criterion (AIC) as well as by performing a Kolmogorov-Smirnov (KS) test. This was done with R packages *fitdistrplus* [35] and *simplexreg* [30]. Besides, for each CpG; the simplex, beta and normal distribution parameters were estimated using maximum likelihood estimation (MLE) using *fitdistrplus* [35], *ZOIP* [27] and *VGAM* [36] R packages.

#### Simulations

The purpose of this second modelling analysis was to fit regressions under a realistic scenario. To that end, we used the parameters estimated in the previous section to generate three data sets, one for each distribution: simplex, beta and normal. These synthetic data sets were constructed using parameters randomly selected from real data estimations containing two balanced groups of samples. In practice, this was generated in two steps for each produced data set: 1) 5% of the simulated DMSs were originated from the same distribution by randomly choosing parameters for each group. 2) The remaining 95% of the data were generated from the specific distribution but fixing the same parameters for all simulated samples. In these settings, only the 5% of synthetic CpGs should be selected as DMSs. Six different scenarios were generated according to the simulated number of samples per group: 3, 5, 10, 30, 100 and 500. Each synthetic data set had 2,000 simulated CpGs each.

After synthetic data were generated, regression models (simplex, inflated simplex, beta, inflated beta, normal and quantile) were fitted for each of the simplex, beta and normal simulated data sets to test the best fit for each distribution. Another popular linear model, *limma*, computed on the logit transformed beta-values (the M-values), was also included in the simulation for comparison purposes. Normal data was restricted to the (0,1) interval for the data to be realistic. R packages *simplexreg, ZOIP, betareg, gamlss* and *quantreg* were used to fit regression models. All model results were adjusted for multiple comparisons with the false discovery rate (FDR, [37]).

Simulations were evaluated in terms of true positives (TP), false positives (FP), true negatives (TN), and false negatives (FN). Three measures were computed out of them: sensitivity, specificity and the Jaccard index. The Jaccard index is defined as the TP/total of real tests, i.e. TP/(TP+FP+FN) [14].

#### Model testing for sex comparison

Finally all assessed regression models were fitted in each data set to test for sex differences. The *limma* approach, applied to the transformed M-values was also included in these analyses since this it is a broadly used method in methylation data analyses. Results were appraised by means of the AIC but also studying the number of DMSs obtained and their genomic position.

## Results

### Data distribution exploration

To see how beta-values distribute in the real data, we first inspected the global profile for all sites of each data set. This presented a high variability in the shape among data sets (Figure 2).

Microarray beta-values were less disperse than sequencing data but they all presented a rough bimodal profile. NGS data showed a different behaviour in RRBS and WGBS, depicting a clear bimodal distribution but with lower intermediate 0-1 values than microarrays. RRBS data showed a preponderant concentration on the tails of the distribution.

In terms of the AIC, simplex was the selected distribution for microarray data sets and RRBS (Figures 3 A-C) whilst beta was selected for WGBS data set (Figure 3 D, Supplementary table 1).

**Figure 3:**
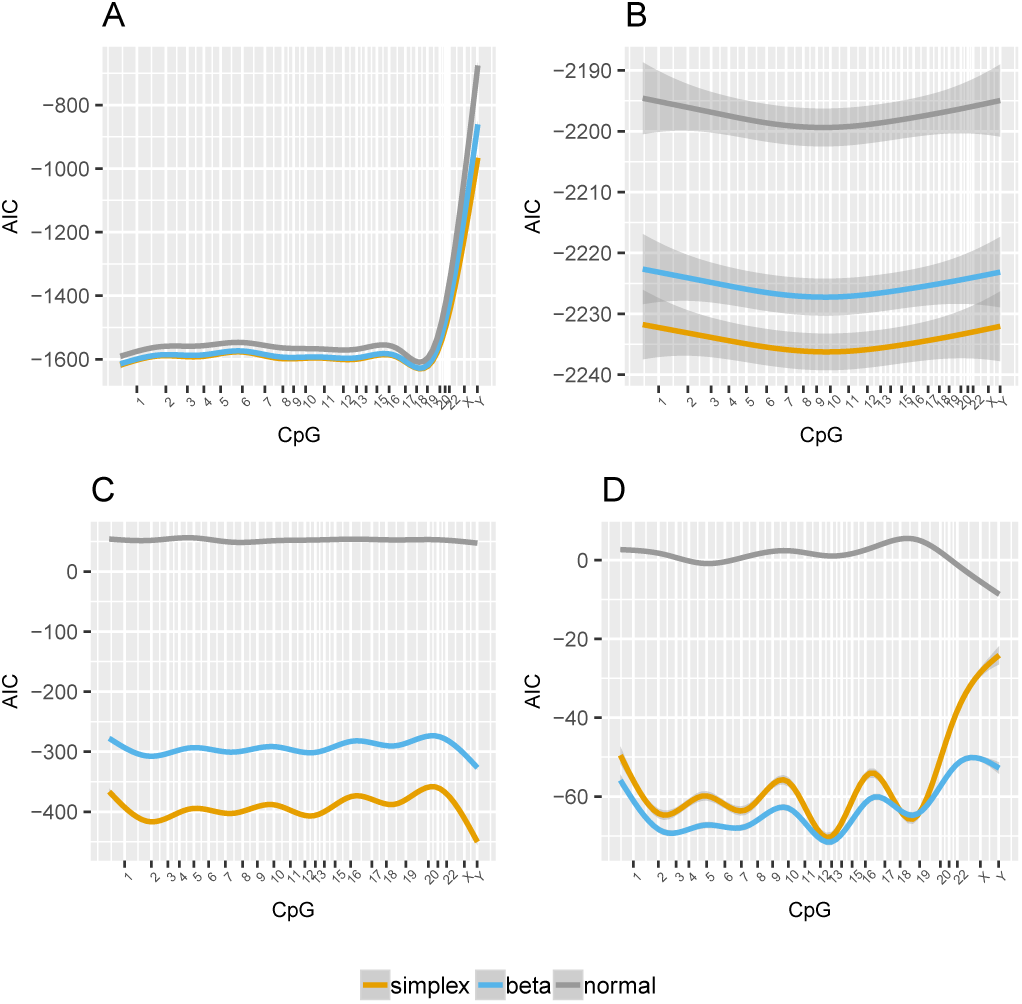
Akaike’s information criterion (AIC) after fitting a simplex, beta or normal distribution for all CpGs in each data set. CpGs are represented in the horizontal axis sorted by chromosomal position. A. array 450k smoking data set. B. array EPIC PBB data set. C. RRBS Ewing sarcoma data set. D. WGBS BLUEPRINT data set.

Simplex, beta and normal distribution parameters were estimated to be used in the simulations. Their distribution was rather disperse in the four data sets (Supplementary figure 3). Of note, similar parameter distributions were found for the two microarray data sets and also for NGS data sets.

### Simulations

Regression models fitted on simulated data (simplex, beta and normal distributed) showed clearly a different behaviour and a better adjustment in the microarray data compared to NGS for the different non inflated models (Figure 4, Table 3, Supplementary figure 4 and Supplementary tables 2-7). For microarray data and according to the Jaccard index, the simplex and beta models were clearly the two best models when data sets were of limited size (n = 3, 5, 10 or 30). Unsurprisingly, as data sets increased in samples, regression models were getting more accurate and the linear model and *limma* showed to be a good alternative. NGS synthetic data modelling was clearly worse in general, with very poor Jaccard indices in general due mainly to an increased FN rate. For larger data sets (n = 100 or 500), in a simplex distribution framework, the *limma* or alternatively, the simplex model, showed the best results being also the normal an alternative for beta or normally distributed NGS data. Inflated beta and simplex models presented overall similar o slightly worse results compared to their corresponding non-inflated models. Of note, the simplex inflated model depicted a higher rate of FP and a lower rate of FN. In addition, quantile regression models underperformed all other models in all situations.

**Table 3:**
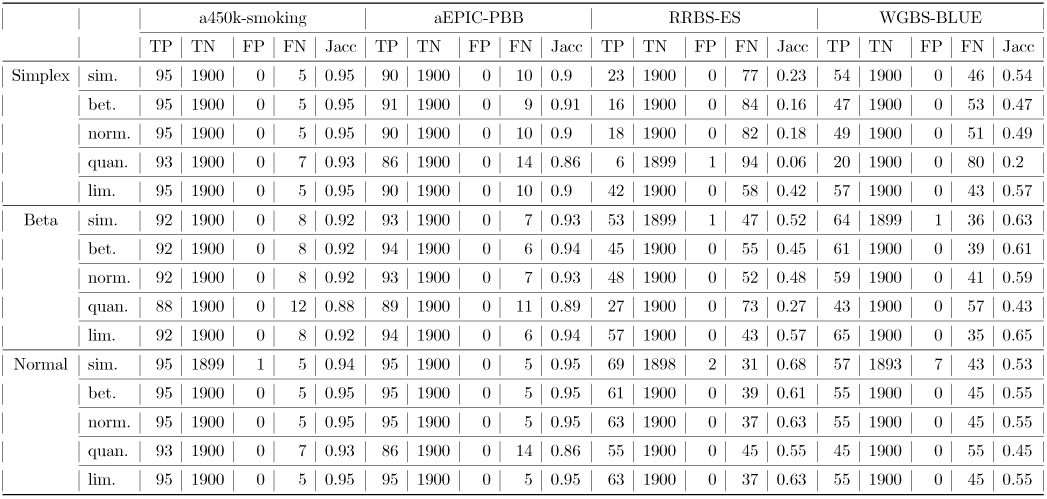
Simulation results. Evaluation measures for synthetic simplex, beta and normal distribution data fitted in to compare two groups of 100 samples through several regression models: simplex (sim.), beta (bet.), normal (norm.), quantile (quan.) and *limma* (lim.). TP: True positives, TN: True negatives, FP: False positives, FN: False negatives, Jacc: Jaccard index. A complete table with all performance measures is to be found in Supplementary table 6.

|  |  | a450k-smoking |  |  |  |  | aEPIC-PBB |  |  |  |  | RRBS-ES |  |  |  |  | WGBS-BLUE |  |  |  |  |
| --- | --- | --- | --- | --- | --- | --- | --- | --- | --- | --- | --- | --- | --- | --- | --- | --- | --- | --- | --- | --- | --- |
|  |  | TP | TN | FP | FN | Jacc | TP | TN | FP | FN | Jacc | TP | TN | FP | FN | Jacc | TP | TN | FP | FN | Jacc |
| Simplex | sim. | 95 | 1900 | 0 | 5 | 0.95 | 90 | 1900 | 0 | 10 | 0.9 | 23 | 1900 | 0 | 77 | 0.23 | 54 | 1900 | 0 | 46 | 0.54 |
|  | bet. | 95 | 1900 | 0 | 5 | 0.95 | 91 | 1900 | 0 | 9 | 0.91 | 16 | 1900 | 0 | 84 | 0.16 | 47 | 1900 | 0 | 53 | 0.47 |
|  | norm. | 95 | 1900 | 0 | 5 | 0.95 | 90 | 1900 | 0 | 10 | 0.9 | 18 | 1900 | 0 | 82 | 0.18 | 49 | 1900 | 0 | 51 | 0.49 |
|  | quan. | 93 | 1900 | 0 | 7 | 0.93 | 86 | 1900 | 0 | 14 | 0.86 | 6 | 1899 | 1 | 94 | 0.06 | 20 | 1900 | 0 | 80 | 0.2 |
|  | lim. | 95 | 1900 | 0 | 5 | 0.95 | 90 | 1900 | 0 | 10 | 0.9 | 42 | 1900 | 0 | 58 | 0.42 | 57 | 1900 | 0 | 43 | 0.57 |
| Beta | sim. | 92 | 1900 | 0 | 8 | 0.92 | 93 | 1900 | 0 | 7 | 0.93 | 53 | 1899 | 1 | 47 | 0.52 | 64 | 1899 | 1 | 36 | 0.63 |
|  | bet. | 92 | 1900 | 0 | 8 | 0.92 | 94 | 1900 | 0 | 6 | 0.94 | 45 | 1900 | 0 | 55 | 0.45 | 61 | 1900 | 0 | 39 | 0.61 |
|  | norm. | 92 | 1900 | 0 | 8 | 0.92 | 93 | 1900 | 0 | 7 | 0.93 | 48 | 1900 | 0 | 52 | 0.48 | 59 | 1900 | 0 | 41 | 0.59 |
|  | quan. | 88 | 1900 | 0 | 12 | 0.88 | 89 | 1900 | 0 | 11 | 0.89 | 27 | 1900 | 0 | 73 | 0.27 | 43 | 1900 | 0 | 57 | 0.43 |
|  | lim. | 92 | 1900 | 0 | 8 | 0.92 | 94 | 1900 | 0 | 6 | 0.94 | 57 | 1900 | 0 | 43 | 0.57 | 65 | 1900 | 0 | 35 | 0.65 |
| Normal | sim. | 95 | 1899 | 1 | 5 | 0.94 | 95 | 1900 | 0 | 5 | 0.95 | 69 | 1898 | 2 | 31 | 0.68 | 57 | 1893 | 7 | 43 | 0.53 |
|  | bet. | 95 | 1900 | 0 | 5 | 0.95 | 95 | 1900 | 0 | 5 | 0.95 | 61 | 1900 | 0 | 39 | 0.61 | 55 | 1900 | 0 | 45 | 0.55 |
|  | norm. | 95 | 1900 | 0 | 5 | 0.95 | 95 | 1900 | 0 | 5 | 0.95 | 63 | 1900 | 0 | 37 | 0.63 | 55 | 1900 | 0 | 45 | 0.55 |
|  | quan. | 93 | 1900 | 0 | 7 | 0.93 | 86 | 1900 | 0 | 14 | 0.86 | 55 | 1900 | 0 | 45 | 0.55 | 45 | 1900 | 0 | 55 | 0.45 |
|  | lim. | 95 | 1900 | 0 | 5 | 0.95 | 95 | 1900 | 0 | 5 | 0.95 | 63 | 1900 | 0 | 37 | 0.63 | 55 | 1900 | 0 | 45 | 0.55 |

**Figure 4:**
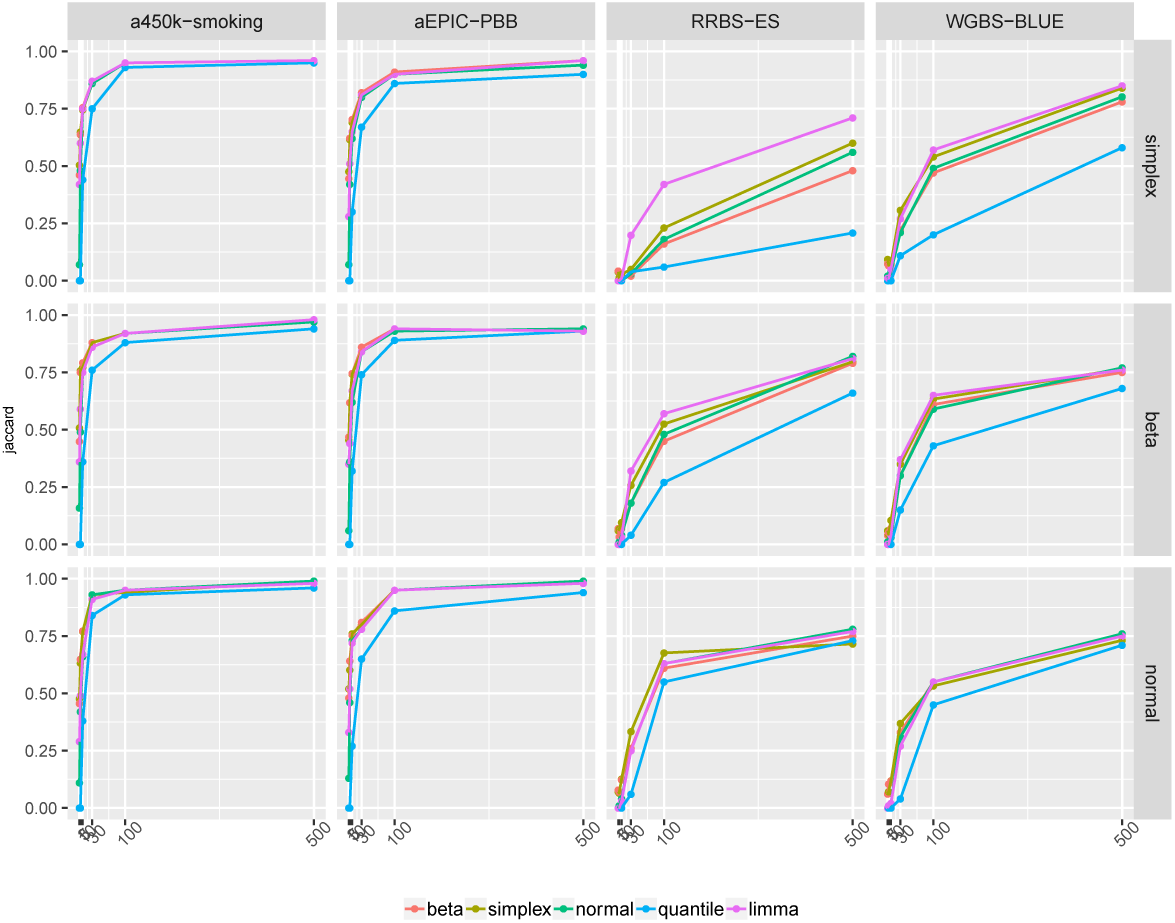
Line plots depicting the Jaccard index for the regression models in the simulations of a two balanced groups comparison with 3, 5, 10, 30, 100 and 500 samples each. Each simulation contains 2000 CpGs with a theoretical 5% of differentially methylated sites. In rows simplex, beta or normal simulated distributions, respectively. In columns the analysis results of each assessed data set: array 450k smoking, array EPIC PBB, RRBS Ewing sarcoma and WGBS BLUEPRINT according to the different regression models, beta, simplex, normal, quantile and limma.

### Sex comparison

The different regression models fitted in the four data sets and evaluated with AIC showed the simplex model to be the most suitable (Figure 5). Results comparing females and males in each data set revealed most of the DMSs to be located in chromosome X although some differentially methylated were also found in autosomal chromosomes (Figure 6, Supplementary table 8). Venn diagrams with the comparison of the results produced by the assessed methods in the four data sets show that non-inflated regression models share in general most of results although simplex leads to more results (Supplementary figure 5). Results for each data set are presented in the following subsections.

**Figure 5:**
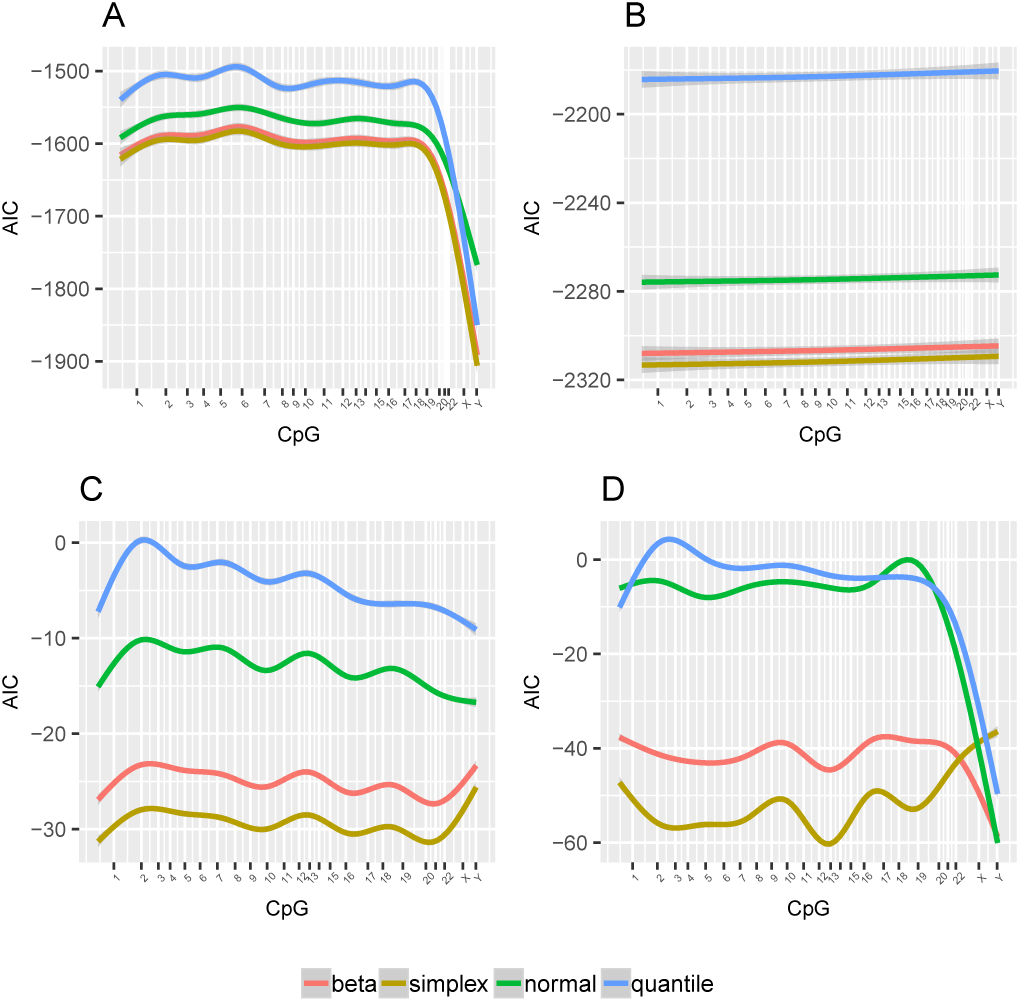
Akaike’s information criterion (AIC) after fitting a beta, simplex, normal or quantile regression model for all CpGs in each data set. CpGs are represented in the horizontal axis sorted by chromosomal position. A. array 450k smoking data set. B. array EPIC PBB data set. C. RRBS Ewing sarcoma data set. D. WGBS BLUEPRINT data set.

**Figure 6:**
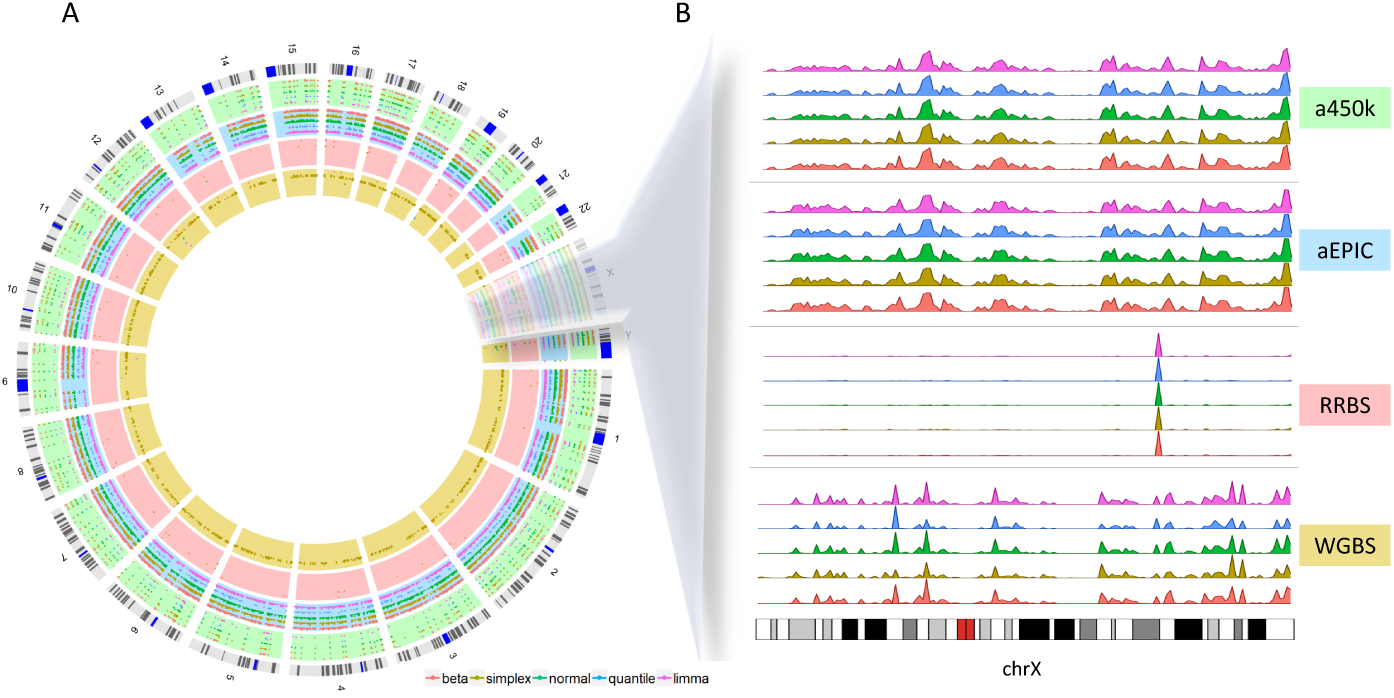
Results of the beta, simplex, normal, quantile and *limma* regression models fitted for all probes in the assessed data sets at an FDR of 1e-08%. A. Circos plot depicting results by chromosome, data set and regression model. B. Density of DMSs obtained by the different regression models located in chromosome X for each assessed data set. In green array 450k smoking data set results, in blue the array EPIC PBB data set results, in red the RRBS Ewing sarcoma data set results and in yellow the WGBS BLUEPRINT data set results.

### array 450k smoking data set

In this smoking data set, 137 females were compared against 327 males. A total of 9,061 different DMSs were found at 5% FDR level for all the assessed models (Supplementary figure 5A), being simplex inflated and non-inflated models the ones presenting the largest number of significant results (12,376 and 11,297 respectively, Supplementary table 8). We observed that between the 66% and 86% of the CpGs that were differentially methylated were located in chromosome X. Beta and normal models detected 1,756 DMSs in autosomes, while simplex only found 281.

### array EPIC PBB data set

Many CpGs were found differentially methylated when comparing 399 females and 280 males at 5% FDR level in the EPIC microarray. The simplex inflated model was again the one with the largest list of DMSs (59,909), while quantile regression detected the lowest number of DMSs (22,523) (Supplementary table 8). We observed a total of 22,492 DMSs shared among all models while 31,102 were found concurrently by the simplex, beta, linear and *limma* models (Supplementary figure 5B). On the other hand, beta and simplex regression had 487 DMSs in common. 317 DMSs were found only by simplex model and 201 by beta model. Chromosome X results represented between 17% and 53% of the total DMSs.

### RRBS Ewing sarcoma data set

Only a few DMSs were obtained in the RRBS data set of Ewing sarcoma samples after comparing 63 females and 96 males. 286 CpGs were selected as DMSs by all methods (Supplementary figure 5C). Of note, 247 DMSs were shared among all models except the quantile model whereas 195 and 36 were found specific of simplex and beta, respectively. Once again, simplex models yielded the largest results, 2,434 and 862 DMSs for inflated and non-inflated respectively (Supplementary table 8). From those, a 59% and 91% belonged to chromosome X, respectively. Notable was the proportion of DMSs found in chromosome X by beta and beta inflated models (94% and 99%, respectively), *limma* (99%) and quantile (100%). Beta-binomial depicted 7,952 DMSs in the chromosome X (99.62%).

### WGBS BLUEPRINT data set

Seventy seven samples had sex information associated, 47 females and 30 males. We found a total of 1,879 DMSs detected by all methods. 3,121 DMSs were found by the simplex, beta, linear and *limma* models and 735 were unique to the beta model (Supplementary figure 5D). Noteworthy, simplex regression model depicted in its non-inflated and inflated versions again 53,637 and 17,923 DMSs (Supplementary table 8), being 11,381 model specific in the non-inflated form. Moreover, simplex models results were spread all over the genome, having a 24% and 46% of the resulting CpGs located in chromosome X, respectively. Beta-binomial results showed 17,129 differentially methylated CpGs, a 61% being located in chromosome X. The remaining models presented a 99% of DMSs in chromosome X.

## Discussion

In this study we have performed comprehensive analyses to gain insights in methylation beta-values distribution, but also to provide practical recommendations in their association to phenotype using accurate regression models. We covered both microarrays and NGS technologies.

Our results showed that real data beta-values matched with a simplex distribution when measured with microarrays or RRBS. In contrast, WGBS beta-values were closer to a beta distribution.

Regression models fitted on simulated yet realistic data presented some interesting results in both microarrays and NGS. Clearly, small data sets with less than 10 samples did not achieve good performance for microarrays nor with less than 30 in NGS. As expected, results were better as the simulated size of groups increased. Microarray simulation results were consistent and for large data sets, simplex, beta but also the linear regression models obtained good evaluation measures. In NGS results were, however, different between RRBS and WGBS. In synthetic data sets of at least 100 samples per group, simplex and *limma* models were the best for WGBS. In RRBS, *limma* was outperforming the other models.

Comparisons between females and males using regression models in the four data sets confirmed on one side, that the simplex regression model could be a good choice and, on the other side, that most of the DMSs were shared among all models except the quantile and located primarily in chromosome X. Some differences were also found in autosomal chromosomes. This is coherent and concordant with previous results [38, 5]. The betabinomial model seemed to capture this overdispersion by presenting more DMSs.

Putting together our results, we have seen that when methylation is assessed with microarrays, beta-values are concordant with a simplex distribution and these can be fitted using a simplex model but also a beta or a linear model, when groups to compare are large. WGBS data set showed to be predominantly beta distributed, but simulations and other analyses performed on this data set showed that a simplex regression model could be used in general whereas *limma* improved the results in some scenarios. RRBS, despite being simplex distributed, presented poor results when adjusted by regression models, with low Jaccard indices influenced by many FN DMSs. However, they could be analyzed using *limma*. Remarkably, quantile regression model is not a good alternative to the other GLMs in the analysis of association with phenotype.

Although we tried to cover many scenarios in our analyses, some limitations need to be mentioned. First of all, the results we provide might be data set dependant but also depend on the R functions and packages used in this study. Besides, there are a few issues that can have a high impact in the data which can condition posterior analyses such as normalization and preprocessing. Pidsley et al. demonstrated for microarrays that with a simple quantile normalization, data performed better than applying other more sophisticated methods [39]. Furthermore, some normalization methods such as quantile normalization lead to non-inflated distributions, with no need to use inflated models. Zero and one inflation can easily overcome by lowering the extremes by a small offset. Sequencing data have also some technical issues such as the over-represented methylation due to higher cycles of PCR [40] or the ‘Spatial correlation’ between the methylation levels of the neighboring [8]. There are other general concerns that need also to be taken into account in the analysis of methylation data such as batch effects [6], tissue heterogeneity, filtering [41], missing values, SNP overlapping and copy number affectation. A careful examination of the data to control these limitations is needed.

## Conclusion

As a final conclusion, we have demonstrated that simplex regression model is a good alternative for the analysis of methylation data whereas the quantile model is not. Results are consistent in microarray data and more heterogeneous in NGS. In summary, this study gives insights into methylation data and analysis, from which our list of recommendations follows: 1) use data sets of at least 10 samples per studied condition for microarrays or 30 in NGS, 2) apply a simplex or beta model in microarray data, 3) apply a linear model in any other case.

## Supporting information

Supplementary file 1. Contains Supplementary figures and Supplementary table 1

## List of abbreviations

AIC: Akaike’s information criterion
DMS: differentially methylated site
FDR: false discovery rate
FN: false negatives
FP: false positives
GLM: generalized linear model
KS: Kolmogorov-Smirnov test
MLE: maximum likelihood estimation
NGS: next generation sequencing
RRBS: reduced representation bisulfite sequencing
sd: standard deviation
SNP: single nucleotide polymorphism
TN: true negatives
TP: true positives
WGBS: whole genome bisulfite sequencing

## Competing interests

The authors declare that they have no competing interests.

## Author’s contributions

JRG and LN had the idea of studying simplex distribution and designed simulation studies. LN wrote the manuscript, carried out simulation studies, performed data analyses and programmed the R functions. JRG helped in writing the final version of the manuscript and oversaw the project. Both authors read and approved the final manuscript.

## Acknowledgements

We thank Júlia Perera and Joan Gibert for carefully reading the manuscript. This work has been supported by the Ministerio de Ciencia, Innovación y Universidades y Fondo Europeo de Desarrollo (RTI2018-100789-B-I00).

