## Supplementary file 1. Contains Supplementary figures and Supplementary table 1 for "Are methylation beta-values simplex distributed?"

### Supplementary Material

August 30, 2019

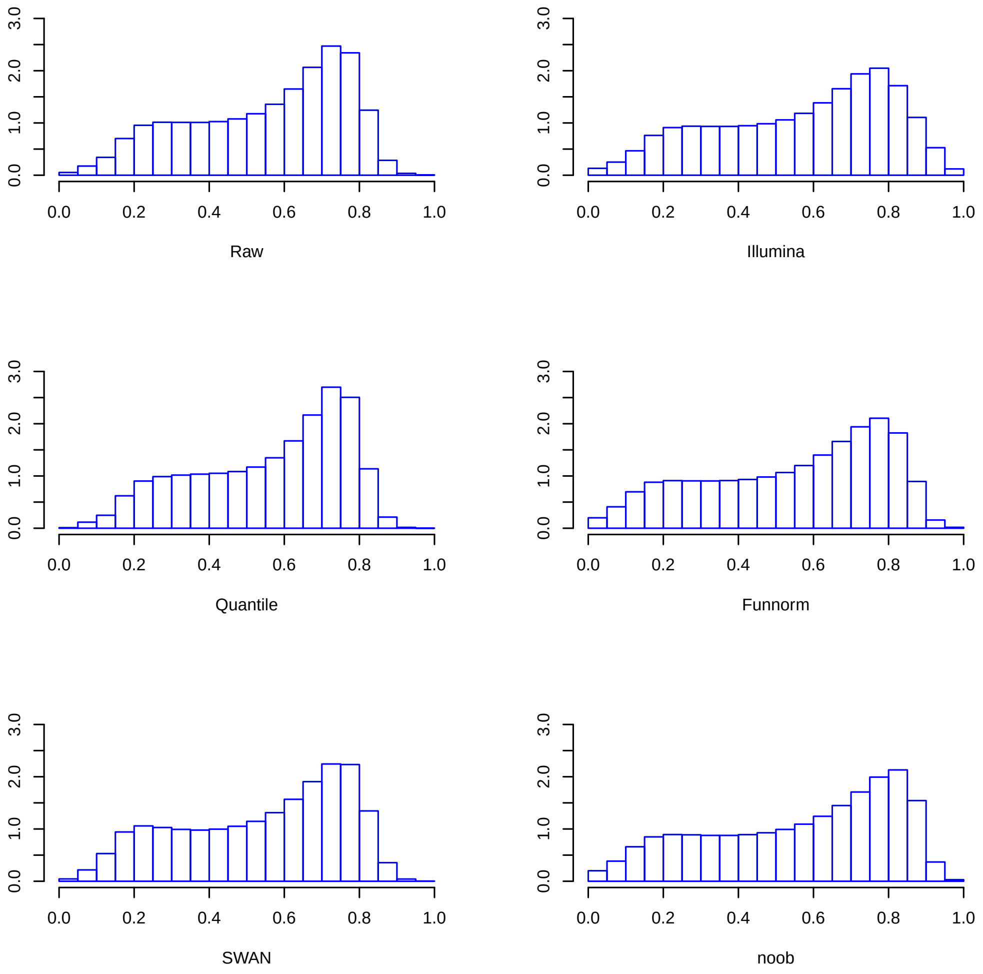

Supplementary figure 1: Global distribution effects of the different normalization methods included in the *minfi* package applied on the aEPIC-PBB data set.

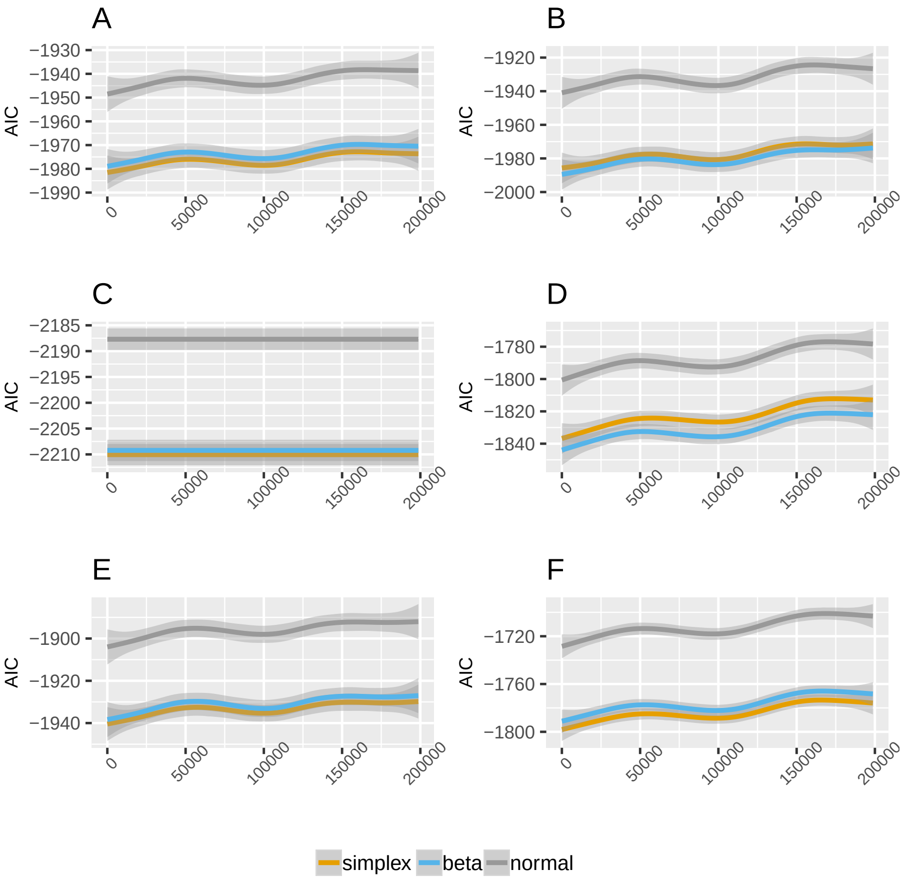

Supplementary figure 2: Akaike's information criterion after adjusting a beta, simplex or normal distribution after the different normalization methods included in the *minfi* package and applied on the aEPIC-PBB data set.

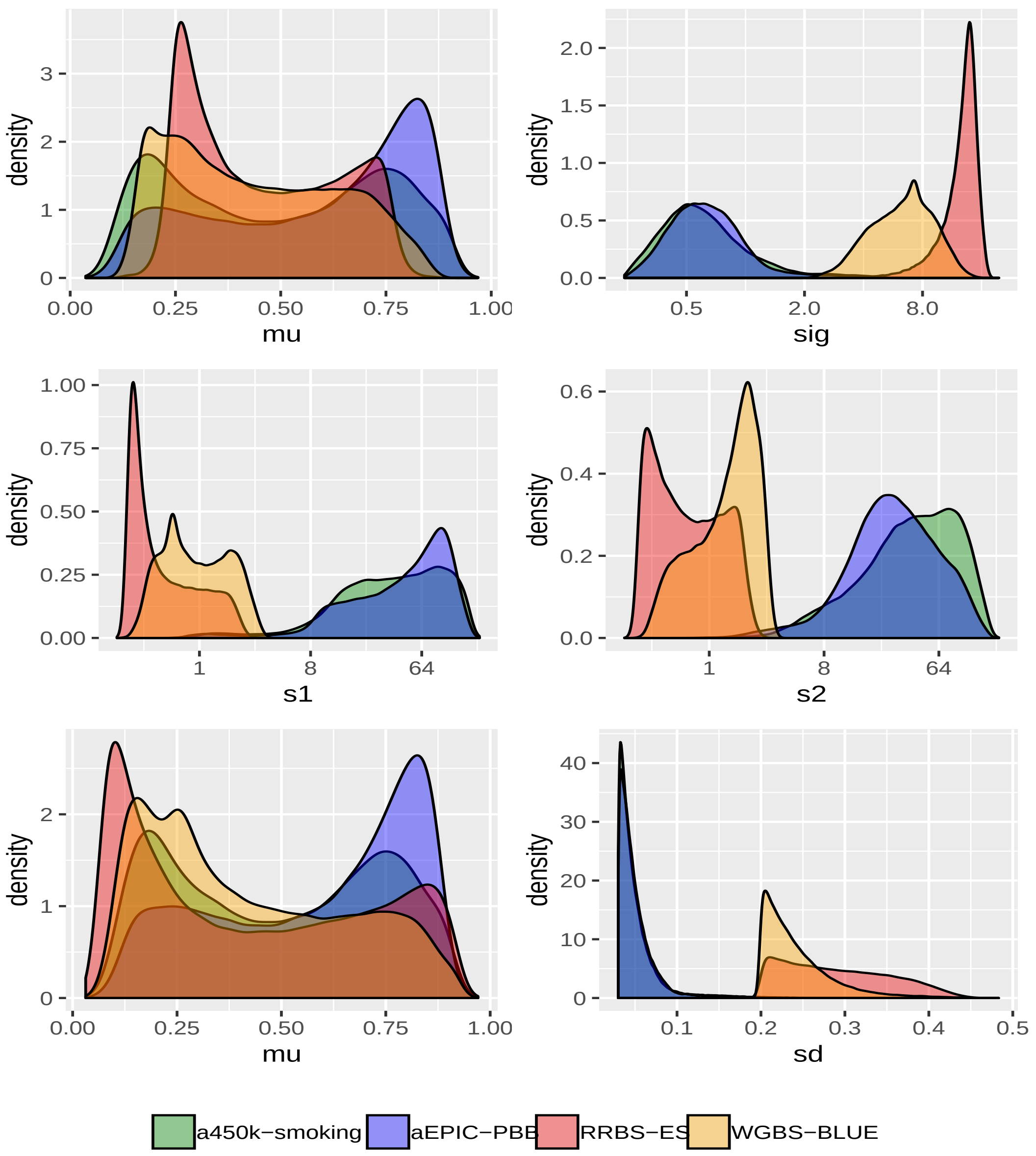

Supplementary figure 3: Parameter estimation distribution obtained by maximum likelihood estimation obtained for parameter in the four analysed data sets. First row corresponds to simplex parameters, second row to beta parameters and third to normal parameters

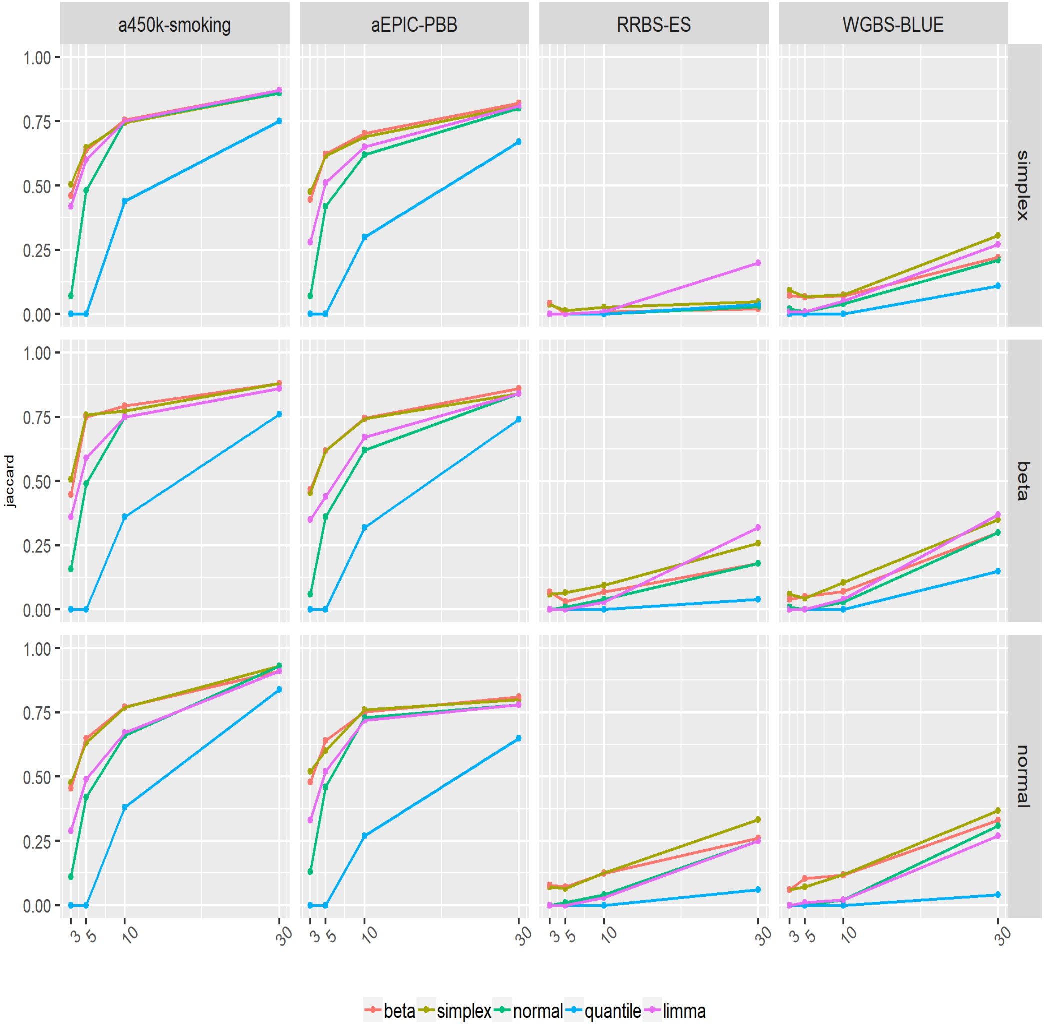

Supplementary figure 4: Line plots depicting the Jaccard index for the regression models in the simulations of a two balanced groups comparison with 3, 5, 10 and 30 samples each. Each simulation contains 2000 CpGs with a theoretical 5% of differentially methylated sites. In rows simplex, beta or normal simulated distributions, respectively. In columns the analysis results of each assessed data set: array 450k smoking , array EPIC PBB, RRBS Ewing sarcoma and WGBS BLUEPRINT according to the different regression models, beta, simplex, normal, quantile and limma.

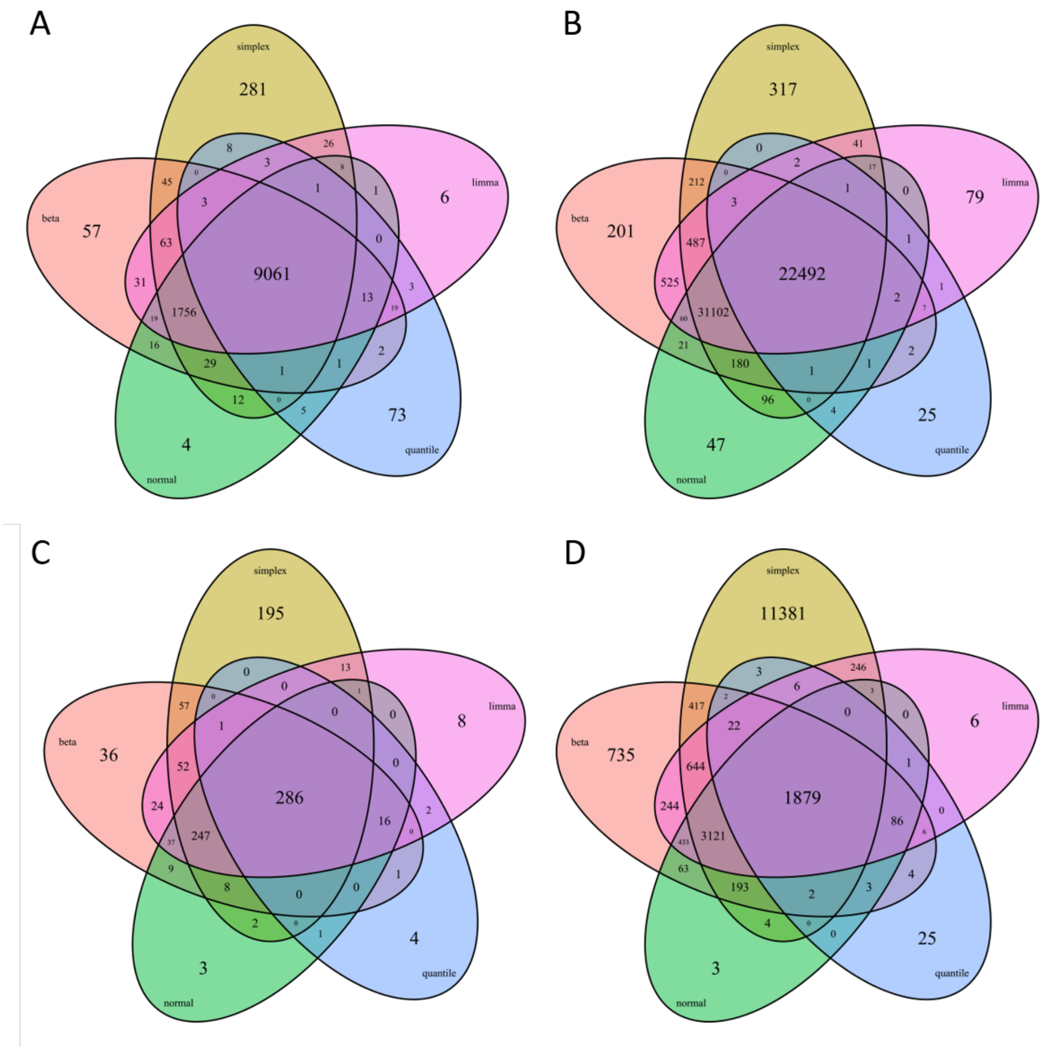

Supplementary figure 5: Venn diagrams comparing results at an FDR of 0.05 after fitting a beta, simplex, normal, quantile and limma models for all probes in the assessed data sets. A. array 450k smoking data set. B. array EPIC PBB data set. C. RRBS Ewing sarcoma data set. D. WGBS BLUEPRINT data set.

Supplementary table 1: Number of CpGs that have simplex, beta or normal distribution as the preferred choice in terms of Akaike's Information Criterion (AIC, the smallest) or the Kolmogorov-Smirnov p-value test (K-S, the highest).

| Data set | Method | simplex | beta | normal |
|---|---|---|---|---|
| a450k-smoking | AIC | 74034 | 19737 | 32179 |
| aEPIC-PBB | AIC | 133548 | 31236 | 33927 |
| RRBS-ES | AIC | 225482 | 43138 | 1949 |
| WGBS-BLUE | AIC | 85606 | 108304 | 10163 |
| a450k-smoking | KS | 74670 | 19028 | 32252 |
| aEPIC-PBB | KS | 120184 | 34779 | 43748 |
| RRBS-ES | KS | 210094 | 18875 | 41600 |
| WGBS-BLUE | KS | 59361 | 79761 | 64951 |

Supplementary table 2: Evaluation measures for synthetic simplex, beta and normal distribution data fitted in to compare two groups of 3 samples by several regression models: simplex (sim.), beta (bet.), normal (norm.), quantile (quan.) and limma (lim.). NAs: number of NAs returned by the model. sens: sensitivity, spec: specificity, TP: True positives, TN: True negatives, FP: False positives, FN: False negatives, jaccard: Jaccard index.

Table is included as a separate excel sheet.

Supplementary table 3: Evaluation measures for synthetic simplex, beta and normal distribution data fitted in to compare two groups of 5 samples by several regression models: simplex (sim.), beta (bet.), normal (norm.), quantile (quan.) and limma (lim.). NAs: number of NAs returned by the model. sens: sensitivity, spec: specificity, TP: True positives, TN: True negatives, FP: False positives, FN: False negatives, jaccard: Jaccard index.

Table is included as a separate excel sheet.

Supplementary table 4: Evaluation measures for synthetic simplex, beta and normal distribution data fitted in to compare two groups of 10 samples by several regression models: simplex (sim.), beta (bet.), normal (norm.), quantile (quan.) and limma (lim.). NAs: number of NAs returned by the model. sens: sensitivity, spec: specificity, TP: True positives, TN: True negatives, FP: False positives, FN: False negatives, jaccard: Jaccard index.

Table is included as a separate excel sheet.

Supplementary table 5: Evaluation measures for synthetic simplex, beta and normal distribution data fitted in to compare two groups of 30 samples by several regression models: simplex (sim.), beta (bet.), normal (norm.), quantile (quan.) and limma (lim.). NAs: number of NAs returned by the model. sens: sensitivity, spec: specificity, TP: True positives, TN: True negatives, FP: False positives, FN: False negatives, jaccard: Jaccard index.

Table is included as a separate excel sheet.

Supplementary table 6: Evaluation measures for synthetic simplex, beta and normal distribution data fitted in to compare two groups of 100 samples by several regression models: simplex (sim.), beta (bet.), normal (norm.), quantile (quan.) and limma (lim.). NAs: number of NAs returned by the model. sens: sensitivity, spec: specificity, TP: True positives, TN: True negatives, FP: False positives, FN: False negatives, jaccard: Jaccard index.

Table is included as a separate excel sheet.

Supplementary table 7: Evaluation measures for synthetic simplex, beta and normal distribution data fitted in to compare two groups of 500 samples by several regression models: simplex (sim.), beta (bet.), normal (norm.), quantile (quan.) and limma (lim.). NAs: number of NAs returned by the model. sens: sensitivity, spec: specificity, TP: True positives, TN: True negatives, FP: False positives, FN: False negatives, jaccard: Jaccard index.

Table is included as a separate excel sheet.

Supplementary table 8: Number of DMSs by chromosome at an FDR of 0.05 after fitting a beta, simplex, normal, quantile and limma models for all probes in the assessed data sets.

Table is included as a separate excel sheet.
